# GNN codon adjacency regulates protein translation

**DOI:** 10.1101/2024.03.16.583757

**Authors:** Joyce Sun, Pete Hwang, Eric D. Sakkas, Yancheng Zhou, Luis Perez, Ishani Dave, Jack B. Kwon, Audrey E. McMahon, Mia Wichman, Mitsu Raval, Kristen Scopino, Daniel Krizanc, Kelly M. Thayer, Michael P. Weir

## Abstract

The central dogma treats the ribosome as a molecular machine that reads one mRNA codon at a time as it adds each amino acid to its growing peptide chain. However, this and previous studies suggest that ribosomes actually perceive pairs of adjacent codons as they take three-nucleotide steps along the mRNA. We examined GNN codons which we find are surprisingly overrepresented in eukaryote protein-coding open reading frames (ORFs), especially immediately after NNU codons. Ribosome profiling experiments in yeast revealed that ribosomes with NNU at their aminoacyl (A) site have particularly elevated densities when NNU is immediately followed (3’) by a GNN codon, indicating slower mRNA threading of the NNU codon from the ribosome’s A to peptidyl (P) sites. Moreover, if the assessment was limited to ribosomes that have only recently arrived at the next codon, by examining 21-nucleotide ribosome footprints (21-nt RFPs), elevated densities were observed for multiple codon classes when followed by GNN. This striking translation slowdown at adjacent 5’-NNN GNN codon pairs is likely mediated in part by the ribosome’s CAR surface which acts as an extension of the A-site tRNA anticodon during ribosome translocation and interacts through hydrogen bonding and pi stacking with the GNN codon. The functional consequences of 5’-NNN GNN codon adjacency are expected to influence the evolution of protein coding sequences.

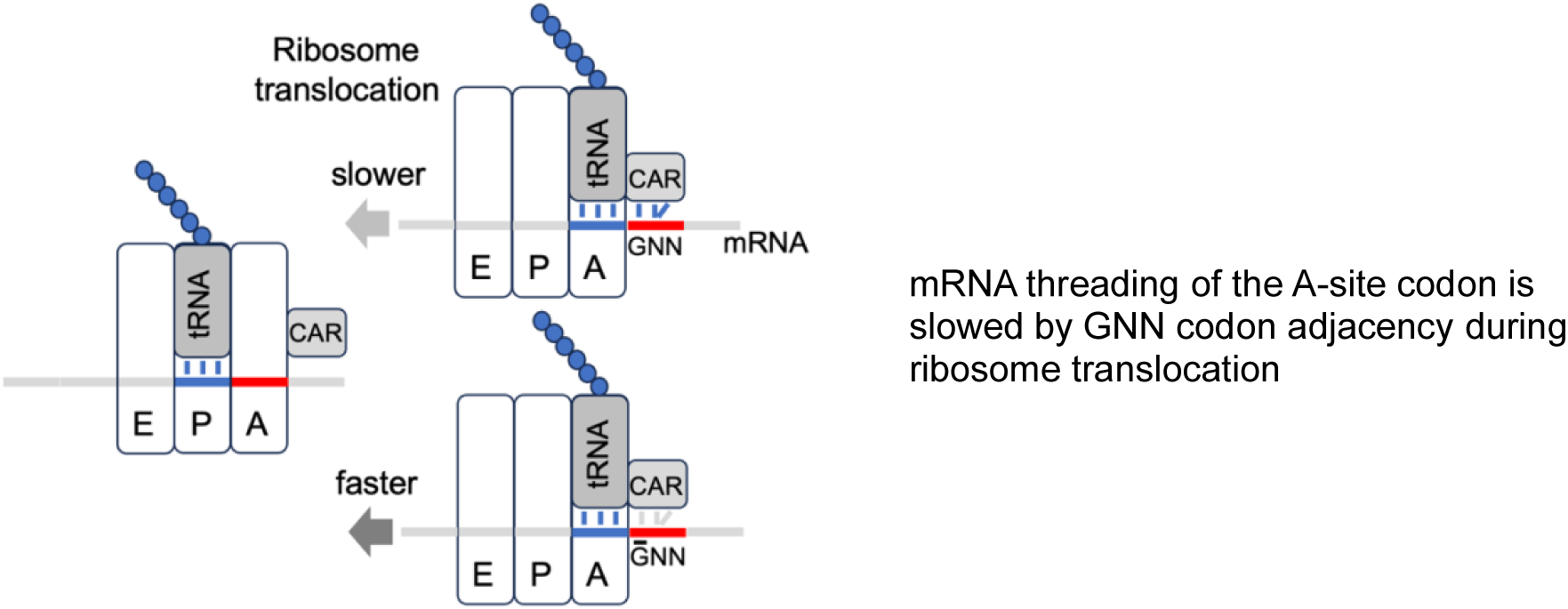
Graphical abstract.

## Introduction

Elevated frequencies of GCN codons (where N = A, C, G or U) have been noted in prokaryote and eukaryote ORFs [1-5] (Fig 1), and their complementarity to sequences in the 530 loop of the ribosome’s 16S rRNA led to speculation that this nucleotide periodicity might be associated with regulation of protein translation [2-5]. With publication of cryo-EM structures of translocating yeast ribosomes, Abeyrathne *et al*. [6] noted hydrogen-bonding between 18S rRNA C1274 and the mRNA nucleotide following (3’ of) the A-site codon. With further examination, we discovered that C1274 is part of CAR [1, 7-9], a three-residue surface that interacts with the full (+1) codon 3’ adjacent to the A-site codon. CAR consists of yeast 18S rRNA C1274 and A1427—which are conserved in prokaryotes and eukaryotes and henceforth referred to as C1054 and A1196, the corresponding 16S rRNA residues in *E. coli*—and R146 of yeast ribosomal protein Rps3 which is only conserved in eukaryotes. The residues of CAR pi stack with each other; C1054 of CAR also pi (base) stacks with nucleotide (nt) 34 of the A-site tRNA anticodon, the partner for the A-site wobble base. This stacking of nt 34 with C1054 of CAR positions CAR as an extension of the anticodon poised to interact with the mRNA +1 codon. Molecular dynamics (MD) simulations of a subsystem of the decoding center neighborhood of translocating ribosomes revealed that the interaction between CAR and the +1 codon is particularly strong during stage II of translocation [9] and the interaction showed a remarkable sequence preference for GNN codons [7-9]. The discovery of CAR and its sequence-specificity for +1 GNN codons further supported the hypothesis that GNN codons might participate in a layer of protein translation regulation. Through context sensitive analysis of published ribosome profiling data, we show here that this is indeed the case. Because CAR is anchored through base stacking to tRNA nt 34, we classified codons/anticodons according to their nt 34 and wobble nucleotide identities and found that NNU A-site codons showed particularly pronounced slowdown in ribosome translocation when immediately followed by a +1 GNN codon.

**Fig 1.**
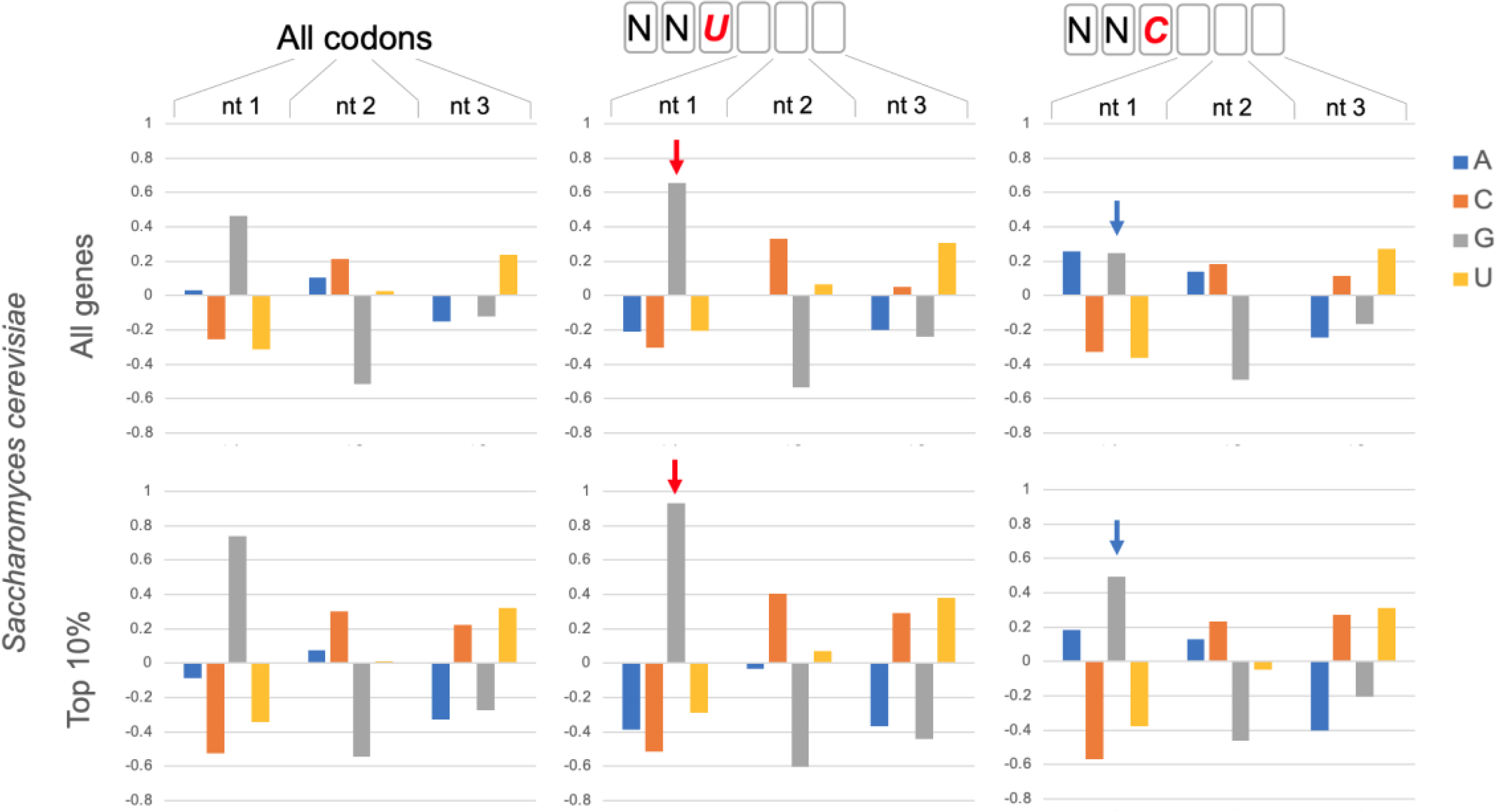
GNN codon adjacency. Codon nucleotide preferences are illustrated by log_2_(f_observed_/f_expected_) using background (expected) frequencies of nucleotides in ORFs regardless of codon position (A:0.325; C: 0.193; G: 0.204; U:0.278). Assessment of all codons in 6703 coding genes in *Saccharomyces cerevisiae* (All codons) showed that G is enhanced in the first nucleotide of codons. This G1 enhancement is significantly more pronounced in codons 3’ adjacent to NNU codons (red arrows; bootstrap p < 0.01) and less pronounced adjacent to NNC codons (blue arrows; bootstrap p < 0.01). The preference for GNN codons is even more pronounced (bootstrap p < 0.01) in genes with high protein expression: the top 10% of protein expressers (Top 10%; 388 genes) from a set of 3868 genes with detectable protein expression in a genomic-scale western and reporter gene analysis in yeast[11].

## Results and Discussion

### NNU codons have distinctive properties

Each NNU codon shares the same tRNA with its corresponding NNC codon containing the same nucleotide identities at positions 1 and 2 of the codon. The anticodons of these tRNAs have either a G34 or I34 (inosine 34) nucleotide, both of which can base pair with the wobble U or C nucleotides of the mRNA A-site codon. I34 (adenine 34 converted to inosine 34) is a eukaryote innovation that is used by NNU/C codon pairs in cases where the corresponding NNA codon also codes for the same amino acid (because inosine can also base pair with wobble A [10]). NNU codons are significantly more abundant than NNC in yeast ORFs (Fig S1A), although for NNU codons that utilize tRNA G34 this trend is less strong in ORFs of genes with high protein expression (Fig S1B, C). Unlike other codon classes, NNU codons are significantly more likely to be immediately followed by a GNN codon (Figs 1 and S2) and this trend is particularly strong for ORFs of genes with high protein expression (Fig 1, Top 10%), suggesting that this codon adjacency may have functional significance. Indeed, position weight matrices for coding sequences of multiple eukaryotes revealed elevated GNN frequencies after NNU codons (Fig S3A), whereas testing of several prokaryote coding sequence showed elevated GNN frequencies after both NNU and NNC codons (Fig S3B).

### GNN adjacency slows protein translation

These hints that NNU GNN codon adjacency might be related to ribosome function led us to examine protein translation rates of NNU codons with and without adjacent GNN codons.

Protein translation rates were assessed by looking at ribosome footprint (RFP) densities on mRNAs which provides a proxy for translation rate where higher density implies slower translation [12] because codons with slower translation receive more footprint “snapshots.”

We analyzed published yeast ribosome profiling data [13] that utilized translation inhibitors tigecycline (TIG) and cycloheximide during harvesting of cell extracts to ensure synchronous cessation of ribosome movement along the mRNA. We examined NNU and NNC codons located at the ribosome A site and compared the ribosome densities for NNU/NNC codon pairs where the codons were followed or not followed by +1 GNN. For all NNU codons (except AUU and CCU, see Table S1), ribosome densities were significantly higher when the codon was adjacent to a 3’ +1 GNN codon compared to when not followed by a +1 GNN codon (bootstrap p < 0.01; blue bars in Fig 2 and replicate data set Fig S3). These trends were observed whether the NNU codon utilized tRNA G34 or I34. Similar consistent trends were not observed for NNC codons (Figs 2 and S3, orange bars; Table S1) or with NNA A-site codons (except GUA codons, Fig S5, Table S1). However, several A-site NNG codons had significantly elevated ribosome densities when followed by +1 GNN and two (AGG and CAG) showed significantly depressed densities (see Fig S5 legend, Fig 3 and Table S1). The pronounced density increases for most NNU codons are consistent with previous observations [14] that codons with wobble U:G base pairing are translated slowly.

**Fig 2.**
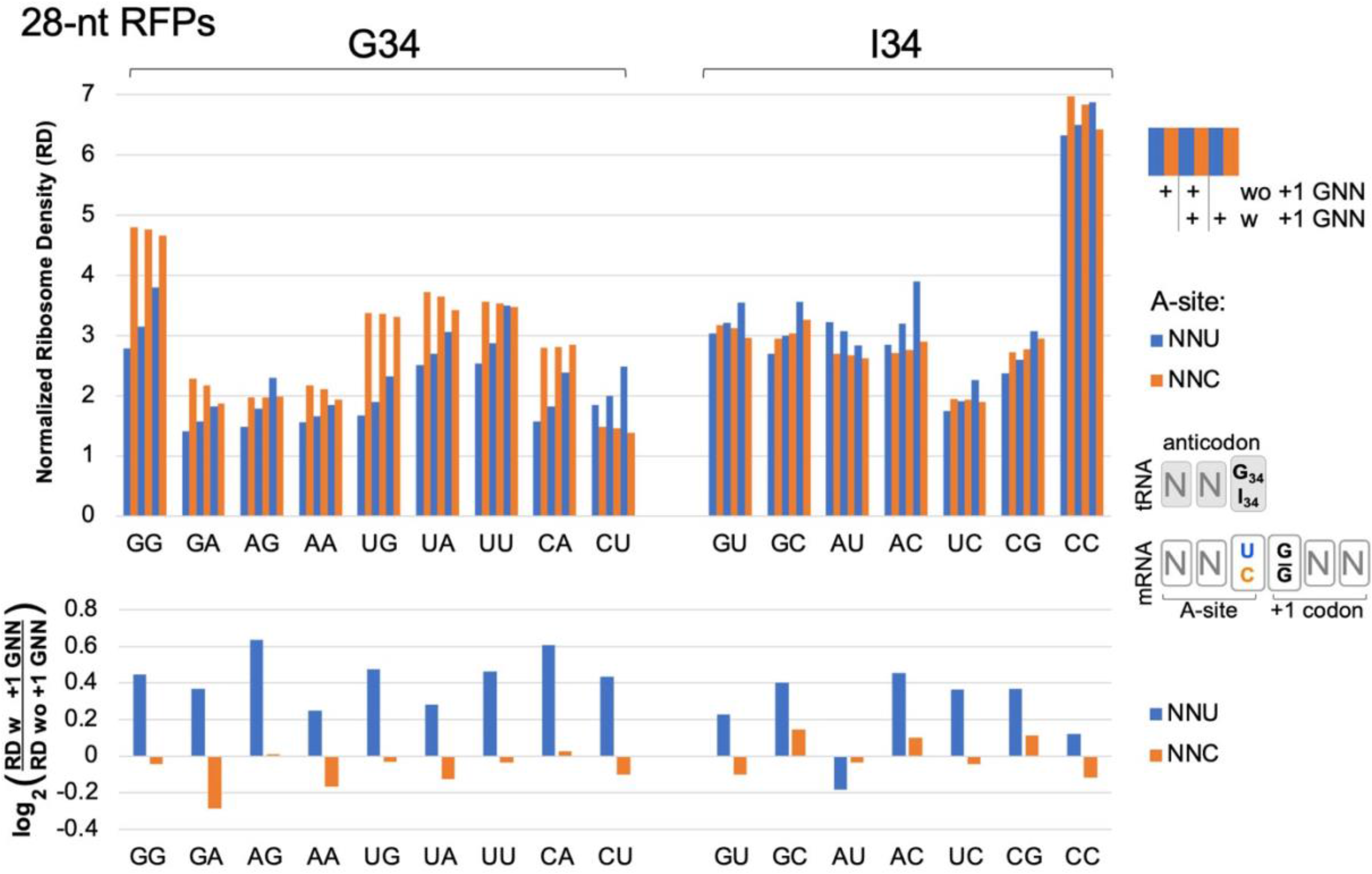
Ribosome densities at A-site NNU/C codons. Analysis of published [13] ribosome profiling data for 629,141 footprints of length 27-30 (28-nt RFPs) in ORFs of 2,132 genes. A-site NNU codons (blue bars) that utilize tRNA G34 or I34 have higher ribosome densities (except AUU, see below), suggesting slower translation rates, when immediately followed (3’) by a codon with G at its first position (with (w) +1 GNN compared to without (wo) +1 GNN). NNC codons (orange) do not show these trends. Codons are labeled on the x-axis according to their first two nucleotides. Codon ribosome densities (RD) were normalized relative to other codons in the same mRNA. Similar results were observed in an independent replicate experiment from the same study [13] (WT2, Fig S4). Bootstrap analysis (2-tailed, p < 0.01) confirmed that in both replicate experiments all NNU codons, except AUU and CCU, had higher densities when followed by +1 GNN (see Table S1). CCU had higher densities in one replicate; AUU had lower densities in one replicate. No NNC codons had consistently higher or lower densities in both replicates.

**Fig 3.**
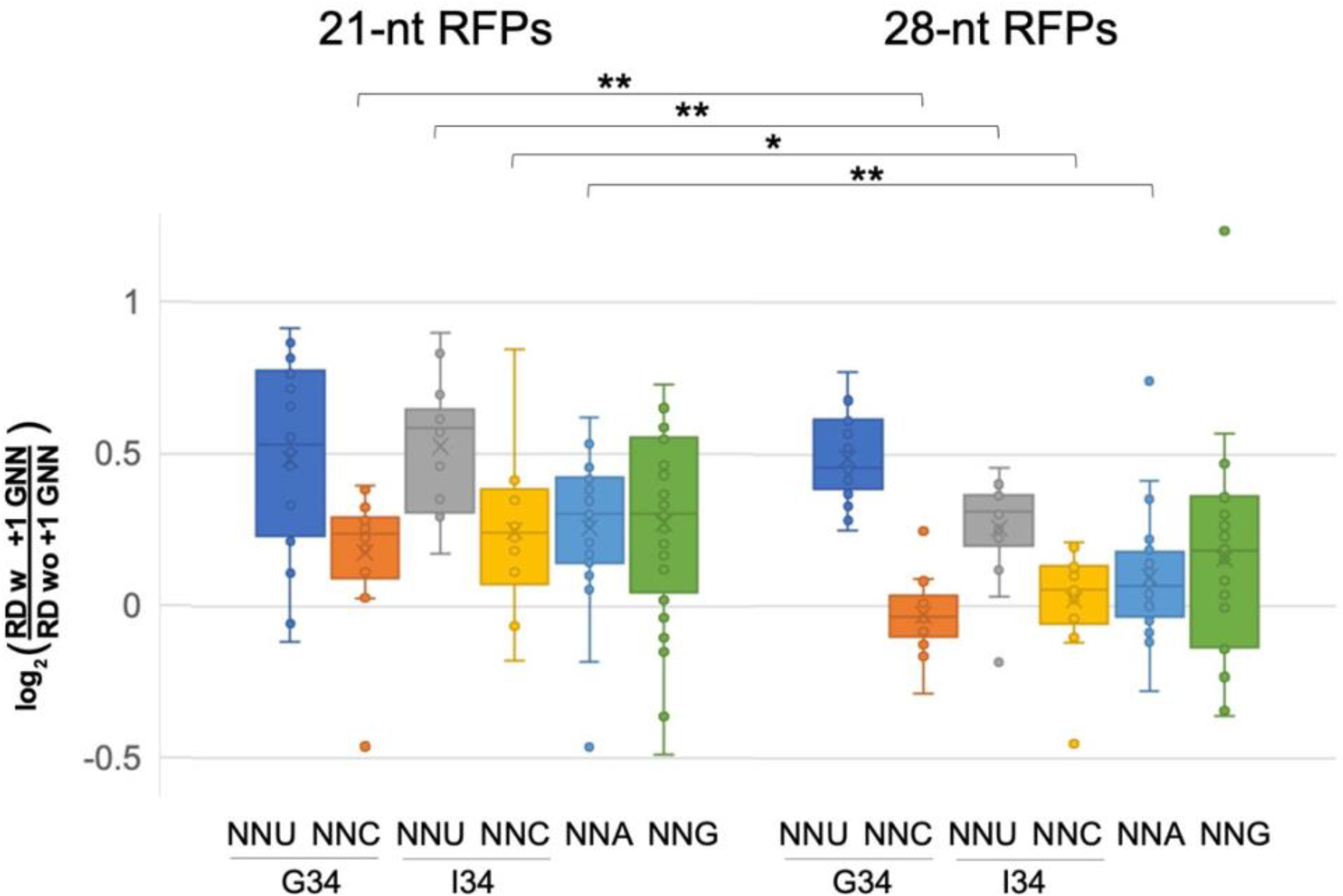
Pre-accommodation and translocation ribosomes are slowed by +1 GNN codons. Analysis of 21-nt ribosome footprints (RFPs) showed that +1 GNN codons led to elevated ribosome densities, for all codon classes, and particularly for A-site NNU codons. 28-nt footprints showed similar trends but +1 GNN codons led to negligible density increases for NNC and NNA codons (see Tables S1 and S2). The density distributions for 21- and 28-nt RFPs were compared using 2-tailed Student’s t-tests (p < 0.05 *; p < 0.01 **): compared to the 21-nt RFPs, +1 GNN codons were associated with lower increases in 28-nt RFP densities for A-site NNC, NNA and NNU (G34). The 21-nt RFPs likely represent pre-accommodation ribosomes that have not accommodated a tRNA at their A-site, whereas the 28-nt RFPs represent translocating ribosomes that have an A- and P-site tRNA. Data was collected from two independent replicate experiments [13] (WT1 and WT2, Figs 2, S4-S7, Tables S1 and S2) and the average normalized ribosome densities for each codon type (one for each replicate) were included in the whisker plot. However, CGA 21-nt RFPs for replicate WT2 were excluded due to very small numbers of CGA GNN codon pairs with above-threshold ribosome densities (>1 footprint per 10 nt).

The above analysis was performed for ribosome footprints of 27-30 nucleotides (28-nt RFPs) which map ribosomes with tRNAs at their P and A sites. Ribosomes also exhibit shorter 20-22 nt footprints (21-nt RFPs) which represent pre-accommodation ribosomes that have not received an A-site tRNA and were frozen in this state because the cell lysates were treated with TIG which blocks tRNA accommodation [13]. Although fewer genes had above-threshold densities of the shorter footprints (>1 footprint per 10 nt), our examination of 21-nt RFPs showed that many classes of codons at the A-site had higher ribosome densities when followed by +1 GNN (bootstrap p < 0.01; Figs 3, S6 and S7, Table S2). Most NNU codons showed high increases in density, including NNU codons that utilize I34 tRNAs (Fig 3, Table S2). Also of note, many NNC and NNA codons, which did not show significant +1 GNN-mediated density increases for 28-nt RFPs, nevertheless had significant density increases for the 21-nt RFPs. These observations suggest that there is a general slowing of pre-accommodation ribosomes when A-site codons are followed by +1 GNN.

### The CAR surface is ideally positioned to mediate +1 GNN regulation

The mechanisms by which pre-accommodation ribosomes (21-nt RFPs) are slowed by +1 GNN codons are unknown. However, once ribosomes have accommodated the correct A-site tRNA (28-nt RFPs), the effects of +1 GNN codons on translation rates are likely mediated by the ribosome CAR surface which is located next to the A-site tRNA anticodon during translocation [1, 7-9]. Supporting this hypothesis, MD analysis of the translocation ribosome decoding center with A-site NNU codons followed by +1 GCU or +1 CGU revealed significantly higher H-bond interactions between CAR and +1 GCU compared to +1 CGU (Fig 4A, B; Student’s t-test p < 0.01). These effects were most pronounced for NNU codons that utilized tRNA G34 and were more muted for those that utilized tRNA I34. A-site NNC codons also showed higher H-bonding between CAR and +1 GCU than +1 CGU, although these effects were less pronounced than for corresponding NNU codons (Fig 4A, B). Our analysis included assessment of the same A-site codon pair (CCU/CCC) tested with tRNA G34 or tRNA I34—the former is utilized by bacterial and archaea tRNAs and the latter by eukaryotes—and this direct comparison of otherwise identical ribosome subsystems confirmed that I34 led to more muted differences between +1 GCU and +1 CGU compared to G34 (Fig 4A, B).

**Fig 4.**
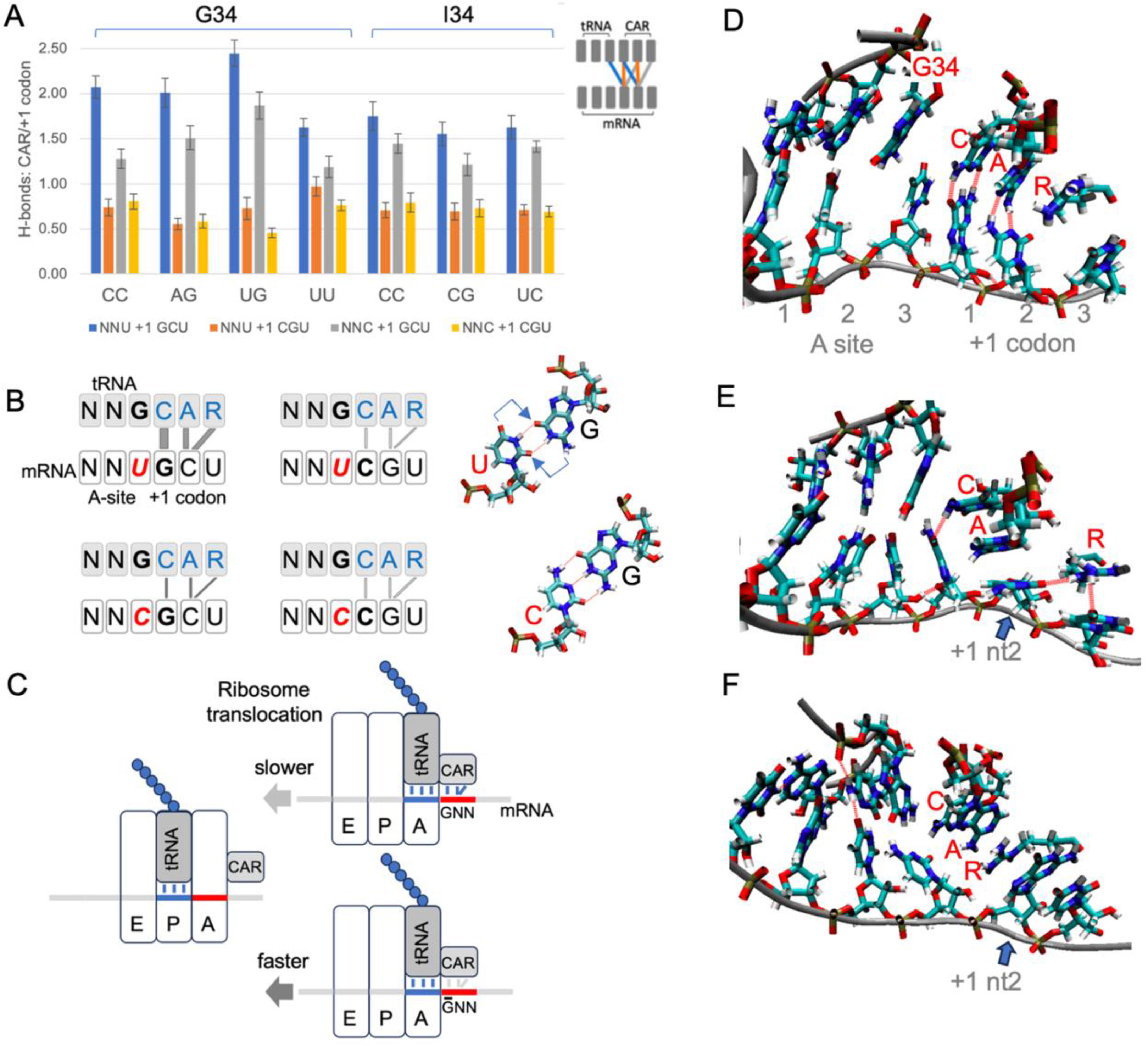
CAR is positioned to mediate +1 codon translation regulation. (A) H-bonding levels between CAR and the +1 codon (see schematic insert and panel B) were assessed in MD trajectories for different combinations of A-site and +1 codons in a subsystem of the translocation stage II ribosome (each construct was tested using 30 MD replicates). The CAR surface preferentially H-bonds in subsystems with +1 GCU codons, and H-bonding is more pronounced with A-site NNU codons that utilize tRNA G34. (B) Schematic summary showing stronger H-bonding between CAR and +1 GCU codons. Wobble U:G base pairing is displaced (arrows) compared to C:G, potentially influencing CAR behavior. (C) Stronger interactions between CAR and the +1 GNN codon are hypothesized to slow threading of the mRNA when the A-site tRNA and its base-paired codon translocate to the ribosome P site. (D) Frame of MD trajectory showing H-bonding between CAR and +1 codon nucleotides 1 and 2 (broken red lines; A-site UUU +1 GCU shown here). (E) In subsystems with +1 GCU, A1196 of CAR often stacks with +1 codon nucleotide 2 (blue arrow; A-site UUU +1 GCU). (F) Subsystems with +1 CGU frequently show stacking of R146 of CAR with +1 codon nucleotide 2 (blue arrow; UGU +1 CGU; also see Fig S8).

Our MD analysis also revealed characteristic pi and pi-cation stacking behaviors of CAR (Fig 4D, E, F, Fig S8). G34 or I34 pi stacked with C1054 of CAR for both A-site NNU and NNC codons with either +1 GCU or +1 CGU. Moreover, I34, a eukaryote innovation, showed significantly more stacking than G34 (Fig S8B; Student’s t-test p < 0.05). We also observed frequent stacking between nucleotide 2 of the +1 codon and either A1196 or R146 of CAR: the cytosine of +1 GCU tended to pi stack with A1196 of CAR (Fig 4E), and the guanine of +1 CGU often cation-pi stacked with R146 of CAR (Fig 4F).

### A sequence-sensitive ribosome braking system

Each time the ribosome takes a translocation step, the A-site tRNA and its base-paired mRNA codon need to be released from the A site to move across to the ribosome P site. Since CAR is part of the ribosome structure that remains behind as the tRNA and mRNA advance (Fig 4C), any modulation of the strength of the interaction between CAR and the +1 codon could influence the speed of release. We speculate that the differences in H-bonding and pi stacking interactions associated with +1 GCU and +1 CGU could lead to differences in the rates of ratcheting movements of the ribosome during translocation. Even if subtle, the differences summed over multiple codon pairs could lead to substantial effects on overall protein translation rates. Clustering analysis of cryoEM images [6] has defined five stages of translocation of the yeast ribosome. The interaction between CAR and NNU +1 codons is strongest at translocation stage II and then gradually decreases as the ratcheting proceeds onwards through stages III to V as the A-site tRNA and its associated mRNA codon translocate across to the P site [9]. Stronger interactions of CAR with +1 GNN codons may act like a transient brake, slightly reducing the probability of CAR “letting go” of the mRNA +1 codon thereby reducing slightly the rate of translocation (Fig 4C).

Adjacency of several combinations of codons has been implicated previously in the regulation of protein translation [15-17] and abnormal structures with adjacent CGA and CCG (or CGA) at the ribosome’s P and A sites respectively have been associated with ribosome stalling [18]. Our new observation involves adjacent codons in the ribosome’s A and +1 sites and applies to a large majority of codons as they arrive at the decoding center A site switching between slower or faster translation depending upon whether or not the +1 codon starts with a G nucleotide. For A-site NNU codons, this effect is particularly strong and extends into the translocation stages of elongation. The discovery of the ribosome’s CAR surface which behaves as an extension of the A-site tRNA anticodon provides a potential structural basis for ribosome slowdown. The striking sequence specificity of the CAR effect, both with respect to the wobble nucleotide/tRNA nt 34 identities as well as the sequences of the A-site and +1 codons, suggests that codon adjacencies may have evolved in ORFs to favor efficient protein translation and reduced probabilities of ribosome-ribosome collisions and resulting stress responses [19]. The potential selective advantages of codon adjacencies for cells in different contexts await future analysis. Indeed, the centrality of tRNA nt 34 in anchoring CAR to the anticodon is of particular interest since this residue can be differentially modified under different growth conditions, including stress [20-23], potentially affecting CAR function.

## Materials and Methods

### Ribosome profiling analysis

Our analysis used published ribosome profiling data [13] which has high-confidence mapping of ribosome A sites to mRNA sequences because upon harvesting of yeast cells, progress of ribosomes along the mRNA was blocked effectively by antibiotics tigecyclin (TIG) and cycloheximide (CHX). TIG blocked recruitment of new tRNAs into the ribosome A site [24], and CHX blocked translocation of the A-site tRNAs to the P-site [25]. Ribosomes that have recruited an A-site tRNA give rise to 27-30 nt footprints (28-nt RFPs) whereas ribosomes that have not yet accommodated a correct tRNA at the A site give rise to shorter 20-22 nt footprints (21-nt RFPs) [13]. The combination of TIG and CHX effectively froze in time the positions of ribosomes on mRNAs, allowing good quantitative assessment of ribosomes positioned on each A-site codon. Ribosome footprint sequences were aligned with mRNA sequences using STAR Aligner [26]. Alignments of 28-nt RFPs for higher-expression genes were improved with the Ribodeblur algorithm [27]. As described previously [12], ribosome profiles show a three-nucleotide periodicity, and the densities at individual codons were computed by taking the major peaks in normalized ribosome counts and adding the normalized counts from adjacent minor peaks on either side—whose intensities were found to correlate strongly with the intensities of the major peaks. Ribosome counts were normalized to the average codon densities for each gene, and the codons in each gene were given equal weighting in our analysis to codons from all other genes. Using WT1 and WT2 replicate data sets [13], our analysis was limited to genes with average footprint densities above 1 footprint per 10 nt (WT1: 2132 genes, WT2: 1279 for 28-nt RFPs; WT1: 1591, WT2: 1100 for 21-nt RFPs). Statistical significance of differences in A-site codon densities with and without +1 GNN codons was assessed with bootstrap analysis using mean ribosome densities from 10,000 samples of 2132, 1279, 1591 or 1100 genes randomly chosen with replacement. The mean density for codons with +1 GNN was compared with the bootstrap distribution of mean densities for codons without +1 GNN.

### Molecular dynamics simulations

MD was performed as described in Dalgarno *et al*. [7]. Briefly, we used a 494-residue subsystem of the ribosome consisting of the decoding center neighborhood. Residues at the periphery of the subsystem were restrained to retain the conformation of stage II of yeast ribosome translocation (PDB ID 5JUP [6]). Nucleotide identities in the A-site codon and anticodon were changed using AMBER’s tLEaP, and 30 independent replicate trajectories (20 × 60 ns; 10 × 100 ns) were performed for each tested version of the subsystem. RMSD analysis confirmed that trajectory behavior had settled by 20 ns and the trajectory frames after 20 ns were analyzed with cpptraj functions to measure H-bonding between the A-site and +1 codons and the anticodon and CAR surface, and to measure base stacking using distances between the centers of geometry of stacked base rings or the guanidinium group of R146 (of CAR). For purine bases, we used the shorter of the distances from its pyrimidine or imidazole ring. Statistical significance was determined with Student’s t-tests of the 30 MD replicates.

### Codon frequency and information theoretic analysis

Analysis of codon frequencies and construction of position weight matrices for codons following reference codons was coded in Python and graphed with Excel and R. Nucleotide frequencies were used to compute position weight matrices (where weight = log_2_(f_observed_/f_expected_)). Codon frequency differences between NNU and NNC codons were tested by bootstrap analysis of gene samples with replacement. Similar bootstrap analysis was performed to assess the differences in weights for adjacent codons with or without G at position 1 of the +1 codon.

## Acknowledgements

We thank David Beveridge, Joe Coolon, Scott Holmes and Amy MacQueen for discussions, and Henk Meij for technical assistance with high-performance computing.

## Data availability

Code and data files are available upon request.

- ORF analysis: code for sequence and information theoretic analysis.
- Ribosome profile analysis: +1 GNN codon analysis and data.
- MD experiments: Restart structure and topology files and code for molecular dynamics experiments.

## Supporting information

Fig S1. NNU/C codon frequencies.

Fig S2. GNN codon adjacency in yeast.

Fig S3. GNN codons are overrepresented in eukaryotes and prokaryotes.

Fig S4. Independent replicate ribosome profile analysis of NNU/C codons.

Fig S5. Ribosome profile analysis of NNG and NNA codons.

Fig S6. Pre-accommodation footprints.

Fig S7. Pre-accommodation footprints.

Fig S8. Stacking behavior of NNU/C codons.

Table S1. 28-nt RFPs bootstrap analysis

Table S2. 21-nt RFPs bootstrap analysis

## References

1. Barr WA, Sheth RB, Kwon J, Cho J, Glickman JW, Hart F, et al. GCN sensitive protein translation in yeast. PloS one. 2020;15:e0233197. PubMed Central PMCID: PMCPMC7500604.

2. Curran JF, Gross BL. Evidence that GHN phase bias does not constitute a framing code. J Mol Biol. 1994;235(1):389–95. doi:10.1016/s0022-2836(05)80046-4. PubMed PMID: 8289262.

3. Lagunez-Otero J, Trifonov EN. mRNA periodical infrastructure complementary to the proof-reading site in the ribosome. Journal of biomolecular structure & dynamics. 1992;10(3):455–64. Epub 1992/12/01. doi:10.1080/07391102.1992.10508662. PubMed PMID: 1492920.

4. Mendoza L, Mondragon M, Lagunez-Otero J. Interaction of the 530 ribosomal site with regions of mRNA. Biosystems. 1998;46(3):293–8. Epub 1998/07/21. PubMed PMID: 9668965.

5. Trifonov EN. Translation framing code and frame-monitoring mechanism as suggested by the analysis of mRNA and 16 S rRNA nucleotide sequences. J Mol Biol. 1987;194(4):643–52. doi:10.1016/0022-2836(87)90241-5. PubMed PMID: 2443708.

6. Abeyrathne PD, Koh CS, Grant T, Grigorieff N, Korostelev AA. Ensemble cryo-EM uncovers inchworm-like translocation of a viral IRES through the ribosome. Elife. 2016;5. doi:10.7554/eLife.14874. PubMed PMID: 27159452; PubMed Central PMCID: PMCPMC4896748.

7. Dalgarno C, Scopino K, Raval M, Nachmanoff C, Sakkas ED, Krizanc D, et al. The CAR-mRNA Interaction Surface Is a Zipper Extension of the Ribosome A Site. Int J Mol Sci. 2022;23(3). Epub 2022/02/16. doi:10.3390/ijms23031417. PubMed PMID: 35163343; PubMed Central PMCID: PMCPMC8835751.

8. Scopino K, Dalgarno C, Nachmanoff C, Krizanc D, Thayer KM, Weir MP. Arginine Methylation Regulates Ribosome CAR Function. Int J Mol Sci. 2021;22:1335. PubMed Central PMCID: PMCPMC7866298.

9. Scopino K, Williams E, Elsayed A, Barr WA, Krizanc D, Thayer KM, et al. A Ribosome Interaction Surface Sensitive to mRNA GCN Periodicity Biomolecules. 2020;10(6). doi:10.3390/biom10060849. PubMed PMID: 32503152; PubMed Central PMCID: PMCPMC7357141.

10. Lei L, Burton ZF. “Superwobbling” and tRNA-34 Wobble and tRNA-37 Anticodon Loop Modifications in Evolution and Devolution of the Genetic Code. Life (Basel). 2022;12(2). Epub 20220208. doi:10.3390/life12020252. PubMed PMID: 35207539; PubMed Central PMCID: PMCPMC8879553.

11. Ghaemmaghami S, Huh WK, Bower K, Howson RW, Belle A, Dephoure N, et al. Global analysis of protein expression in yeast. Nature. 2003;425(6959):737–41. PubMed PMID: 14562106.

12. Ingolia NT, Ghaemmaghami S, Newman JR, Weissman JS. Genome-wide analysis in vivo of translation with nucleotide resolution using ribosome profiling. Science. 2009;324(5924):218–23. Epub 2009/02/14. doi:1168978[pii]10.1126/science.1168978. PubMed PMID: 19213877; PubMed Central PMCID: PMCPMC2746483.

13. Wu CC, Zinshteyn B, Wehner KA, Green R. High-Resolution Ribosome Profiling Defines Discrete Ribosome Elongation States and Translational Regulation during Cellular Stress. Mol Cell. 2019;73(5):959–70 e5. Epub 2019/01/29. doi:10.1016/j.molcel.2018.12.009. PubMed PMID: 30686592; PubMed Central PMCID: PMCPMC6411040.

14. Stadler M, Fire A. Wobble base-pairing slows in vivo translation elongation in metazoans. RNA. 2011;17(12):2063–73. doi:10.1261/rna.02890211. PubMed PMID: 22045228; PubMed Central PMCID: PMCPMC3222120.

15. Gamble CE, Brule CE, Dean KM, Fields S, Grayhack EJ. Adjacent Codons Act in Concert to Modulate Translation Efficiency in Yeast. Cell. 2016;166(3):679–90. Epub 2016/07/05. doi:10.1016/j.cell.2016.05.070. PubMed PMID: 27374328; PubMed Central PMCID: PMCPMC4967012.

16. Letzring DP, Dean KM, Grayhack EJ. Control of translation efficiency in yeast by codon-anticodon interactions. Rna. 2010;16(12):2516–28. Epub 2010/10/26. doi:10.1261/rna.2411710. PubMed PMID: 20971810; PubMed Central PMCID: PMCPMC2995412.

17. Lin Y, May GE, Kready H, Nazzaro L, Mao M, Spealman P, et al. Impacts of uORF codon identity and position on translation regulation. Nucleic Acids Res. 2019;47(17):9358–67. doi:10.1093/nar/gkz681. PubMed PMID: 31392980; PubMed Central PMCID: PMCPMC6755093.

18. Tesina P, Lessen LN, Buschauer R, Cheng J, Wu CC, Berninghausen O, et al. Molecular mechanism of translational stalling by inhibitory codon combinations and poly(A) tracts. EMBO J. 2020;39(3):e103365. Epub 20191220. doi:10.15252/embj.2019103365. PubMed PMID: 31858614; PubMed Central PMCID: PMCPMC6996574.

19. Wu CC, Peterson A, Zinshteyn B, Regot S, Green R. Ribosome Collisions Trigger General Stress Responses to Regulate Cell Fate. Cell. 2020;182(2):404–16 e14. Epub 20200630. doi:10.1016/j.cell.2020.06.006. PubMed PMID: 32610081; PubMed Central PMCID: PMCPMC7384957.

20. Chan CT, Dyavaiah M, DeMott MS, Taghizadeh K, Dedon PC, Begley TJ. A quantitative systems approach reveals dynamic control of tRNA modifications during cellular stress. PLoS genetics. 2010;6(12):e1001247. doi:10.1371/journal.pgen.1001247. PubMed PMID: 21187895; PubMed Central PMCID: PMCPMC3002981.

21. Endres L, Rose RE, Doyle F, Rahn T, Lee B, Seaman J, et al. 2’-O-ribose methylation of transfer RNA promotes recovery from oxidative stress in Saccharomyces cerevisiae. PloS one. 2020;15(2):e0229103. doi:10.1371/journal.pone.0229103. PubMed PMID: 32053677; PubMed Central PMCID: PMCPMC7018073.

22. Gu C, Begley TJ, Dedon PC. tRNA modifications regulate translation during cellular stress. FEBS Lett. 2014;588(23):4287–96. doi:10.1016/j.febslet.2014.09.038. PubMed PMID: 25304425; PubMed Central PMCID: PMCPMC4403629.

23. Huber SM, Leonardi A, Dedon PC, Begley TJ. The Versatile Roles of the tRNA Epitranscriptome during Cellular Responses to Toxic Exposures and Environmental Stress. Toxics. 2019;7(1). doi:10.3390/toxics7010017. PubMed PMID: 30934574; PubMed Central PMCID: PMCPMC6468425.

24. Jenner L, Starosta AL, Terry DS, Mikolajka A, Filonava L, Yusupov M, et al. Structural basis for potent inhibitory activity of the antibiotic tigecycline during protein synthesis. Proc Natl Acad Sci U S A. 2013;110(10):3812–6. Epub 2013/02/23. doi:10.1073/pnas.1216691110. PubMed PMID: 23431179; PubMed Central PMCID: PMCPMC3593886.

25. Budkevich T, Giesebrecht J, Altman RB, Munro JB, Mielke T, Nierhaus KH, et al. Structure and dynamics of the mammalian ribosomal pretranslocation complex. Mol Cell. 2011;44(2):214–24. doi:10.1016/j.molcel.2011.07.040. PubMed PMID: 22017870; PubMed Central PMCID: PMCPMC3242006.

26. Dobin A, Davis CA, Schlesinger F, Drenkow J, Zaleski C, Jha S, et al. STAR: ultrafast universal RNA-seq aligner. Bioinformatics. 2013;29(1):15–21. Epub 20121025. doi:10.1093/bioinformatics/bts635. PubMed PMID: 23104886; PubMed Central PMCID: PMCPMC3530905.

27. Wang H, Kingsford C, McManus CJ. Using the Ribodeblur pipeline to recover A-sites from yeast ribosome profiling data. Methods. 2018;137:67–70. Epub 2018/01/14. doi:10.1016/j.ymeth.2018.01.002. PubMed PMID: 29330118; PubMed Central PMCID: PMCPMC6261449.

