## Supplemental Figures and Tables for "GNN codon adjacency regulates protein translation"

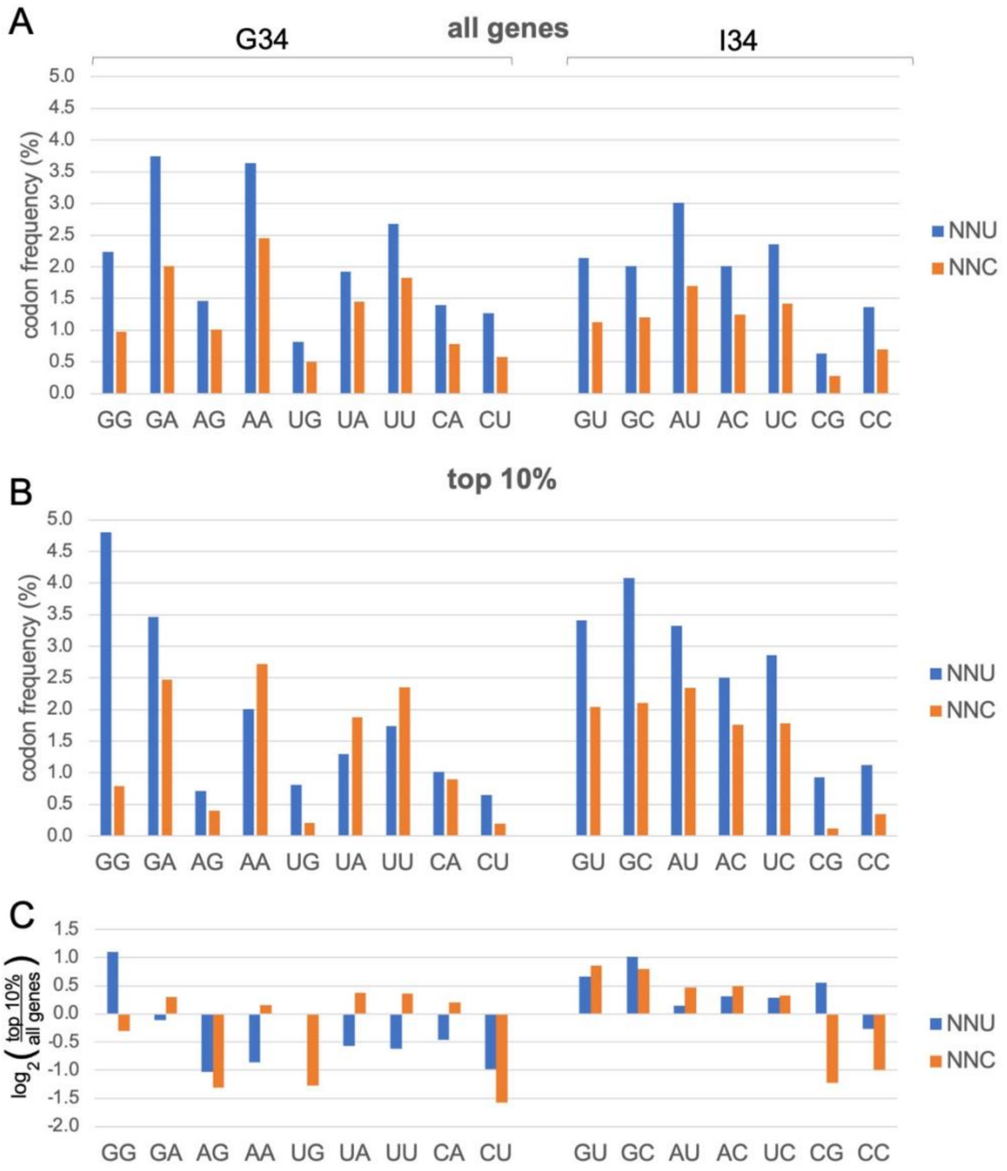


**Fig S1. NNU/C codon frequencies.** (A) For all NNU/C codon pairs, NNU codons have significantly higher frequencies in yeast ORFs than NNC (bootstrap $p < 0.01$). Codons are labeled on the x-axis according to their first two nucleotides. (B) Genes with high protein expression (top 10%, see below) show similar trends. NNU codons have higher frequencies than NNC for all NNU/C codon pairs that utilize tRNA I34 anticodons, and most NNU/C codon pairs that utilize G34, except for AAU/C, UAU/C and UUU/C where NNC codons have significantly higher frequencies (bootstrap $p < 0.01$). We extracted the top 10% of protein expressers from a set of 3868 genes with detectable protein expression in a genomic-scale western and reporter gene analysis in yeast [1]. (C) The NNU/C codon frequencies in high expression genes compared to all genes are significantly elevated for all codons that utilize I34 except CGC, CCC and CCU. In contrast, for codons that utilize G34, most NNU and some NNC codon frequencies are significantly depressed.

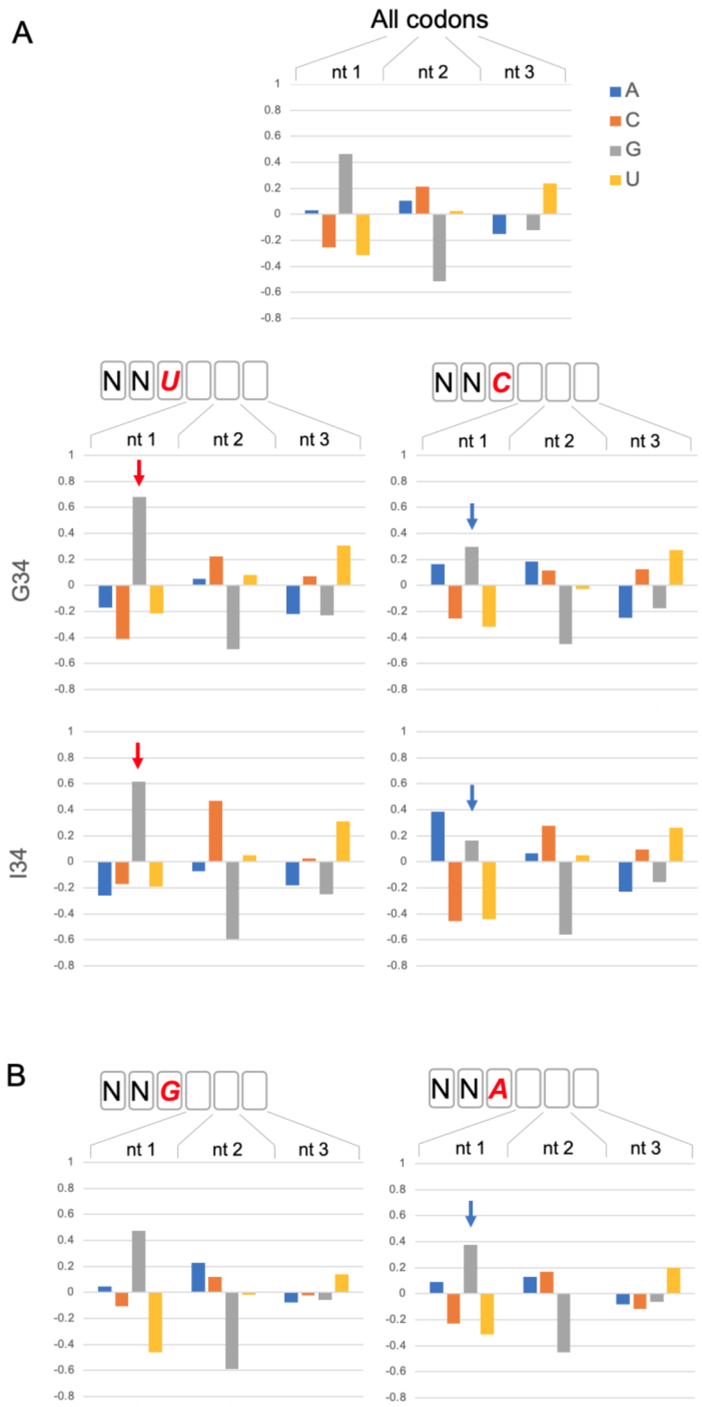


**Fig S2. GNN codon adjacency in yeast.** (A) Position weight matrices of yeast ORFs (6703 genes) show that codons following (3') NNU and NNC have elevated (red arrows) and depressed (blue arrows) GNN frequencies respectively, whether they utilize G34 or I34 tRNAs (bootstrap $p < 0.01$). (B) NNG codons do not show these trends; the codons following NNA have depressed GNN weights (bootstrap $p < 0.01$). Codon nucleotide preferences are illustrated by $\log_2(f_{obs}/f_{exp})$ using background (expected) frequencies of nucleotides in ORFs regardless of codon position (A:0.325; C: 0.193; G: 0.204; U:0.278).

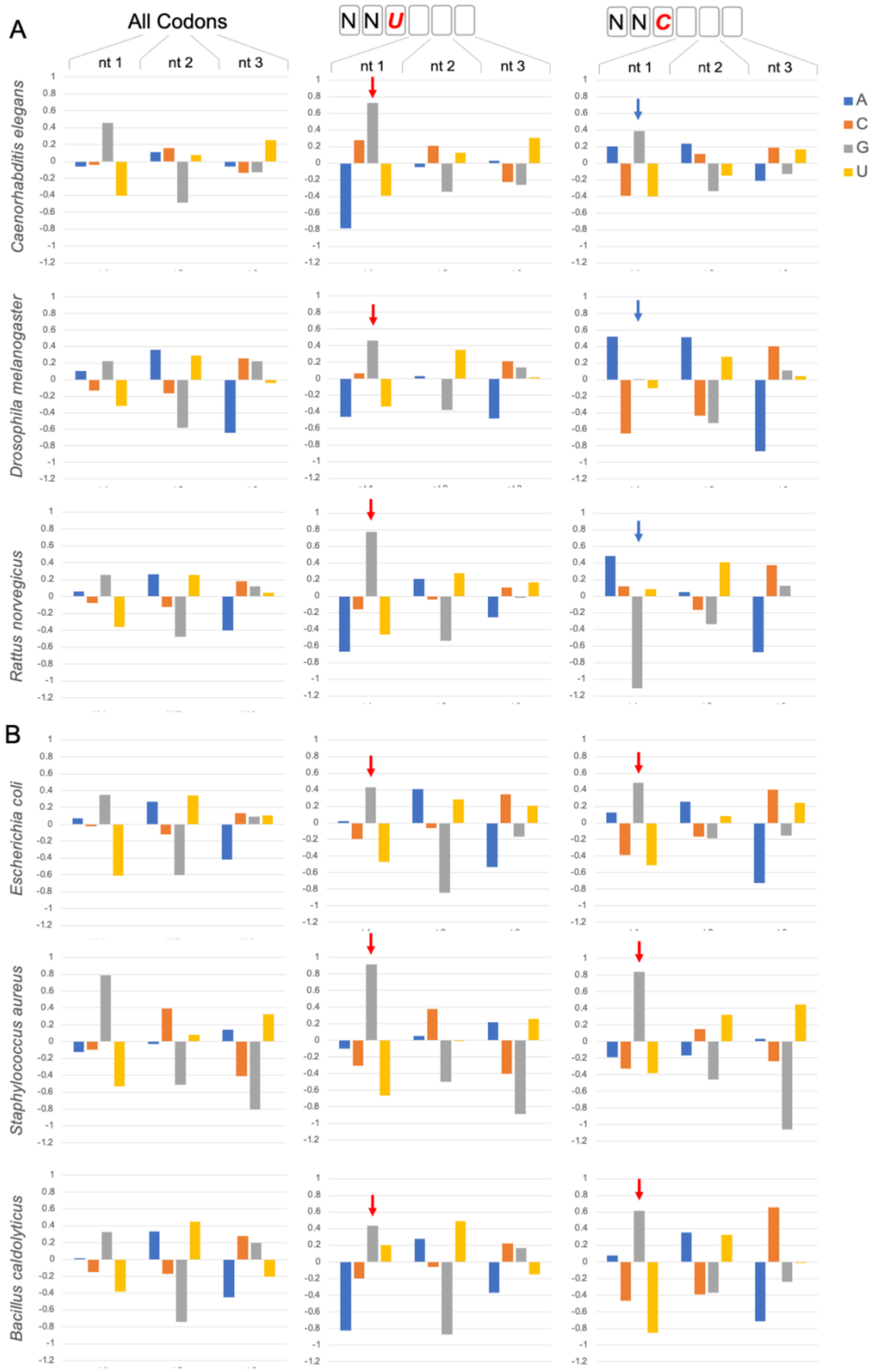


**Fig S3. GNN codons are overrepresented in eukaryotes and prokaryotes.** Position weight matrices illustrating $\log_2(f_{obs}/f_{exp})$ show that GNN codons are overrepresented in ORFs across multiple species. In the eukaryotes (A), GNN is significantly more abundant when 3' adjacent to NNU codons (bootstrap $p < 0.01$, red arrows) and GNN is significantly less abundant when adjacent to NNC codons (bootstrap $p < 0.01$, blue arrows). However, in the prokaryotes (B), GNN is significantly more abundant adjacent to NNU and NNC codons (bootstrap $p < 0.01$, red arrows).

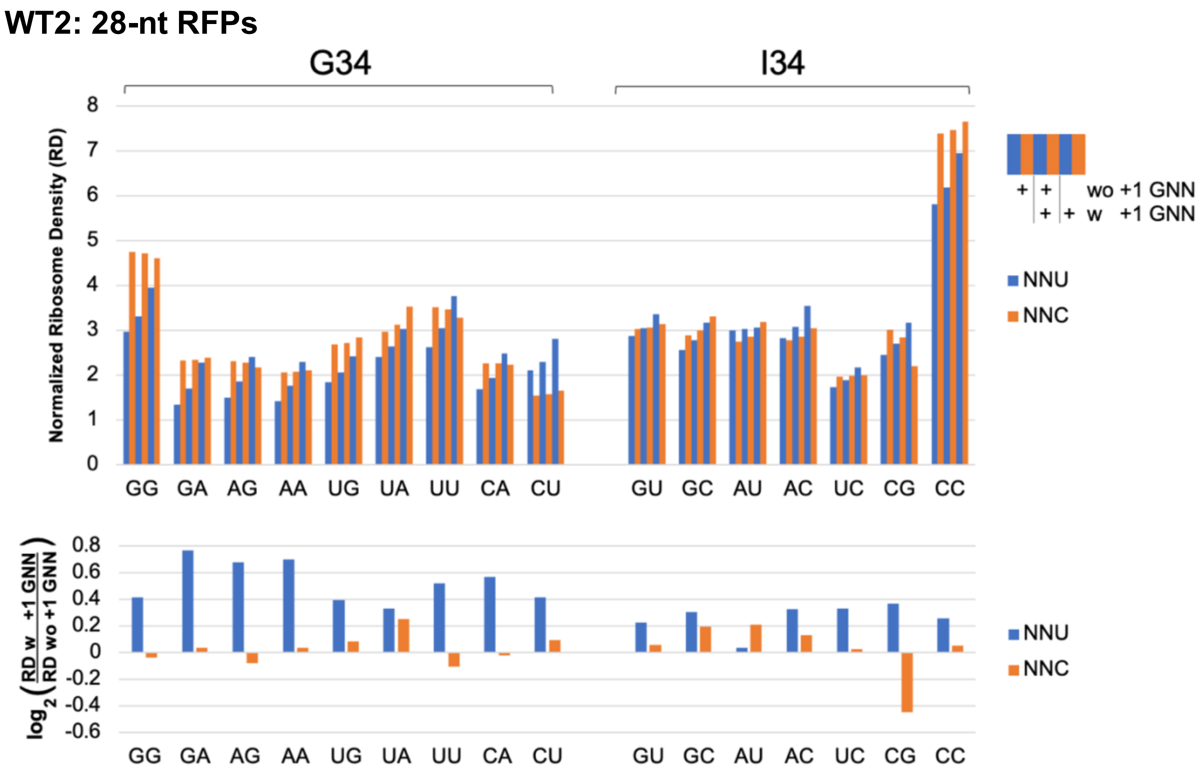


**Fig S4. Independent replicate ribosome profile analysis of NNU/C codons.** Equivalent analysis to Fig 2 but with independent (WT2) replicate dataset from the same study [2].

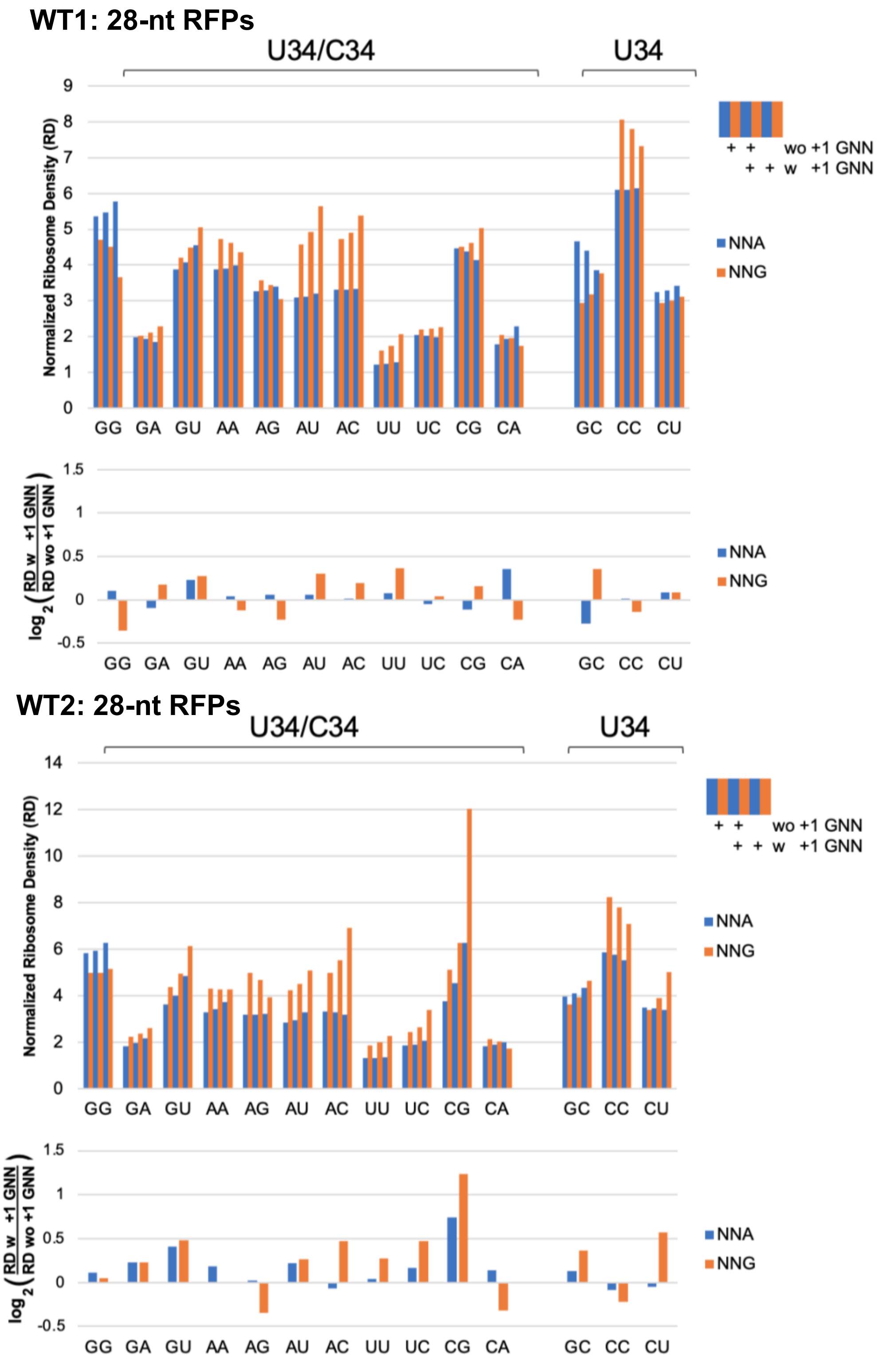


**Fig S5. Ribosome profile analysis of NNG and NNA codons.** Several A-site NNG (GCG, GAG, GUG, AUG and UUG) ribosome profiles (28-nt RFPs) had significantly increased densities when followed by +1 GNN in both replicate experiments [2] (WT1 and WT2; bootstrap $p < 0.01$; Table S1) whereas CAG and AGG had reduced densities. Only one NNA codon (GUA) had significantly increased densities with +1 GNN in both replicates. Data in Figs S5, S6, and S7 is graphed as in Figs 2 and S4. GGG/A, GAG/A and GUG/A codon pairs share the same U34 tRNA; the other NNG/A codon pairs have separate tRNAs for each codon.

**WT1: 21-nt RFPs**

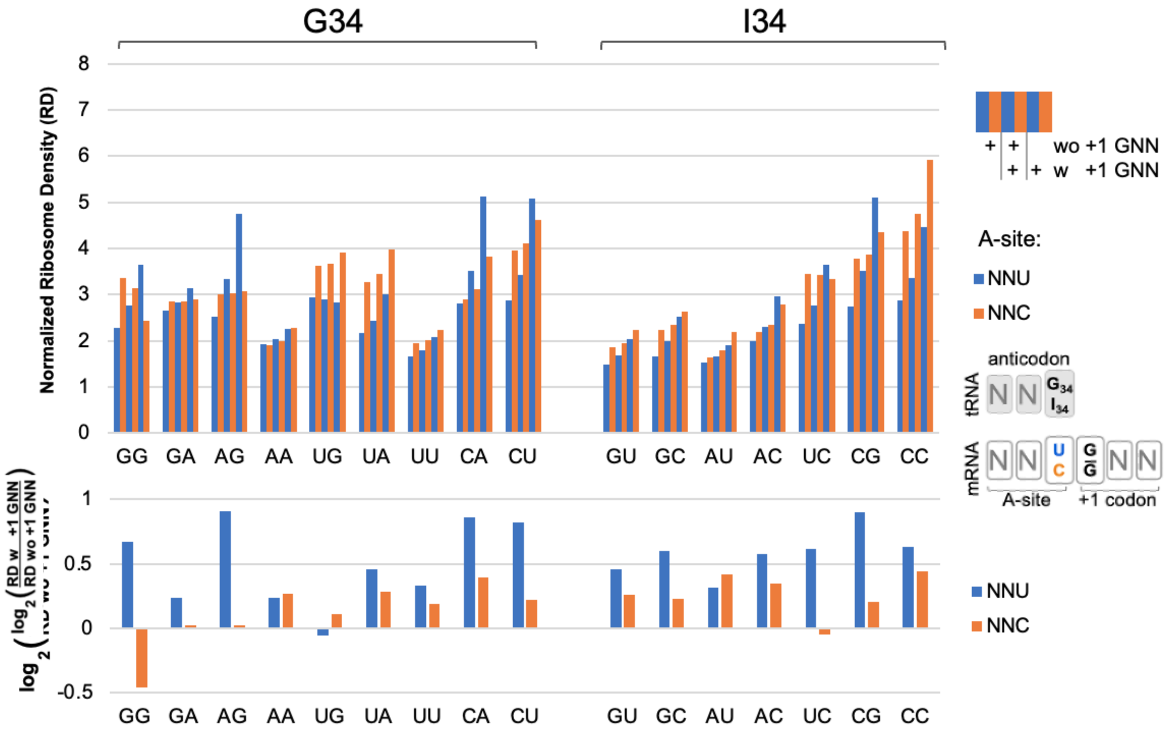


**WT2: 21-nt RFPs**

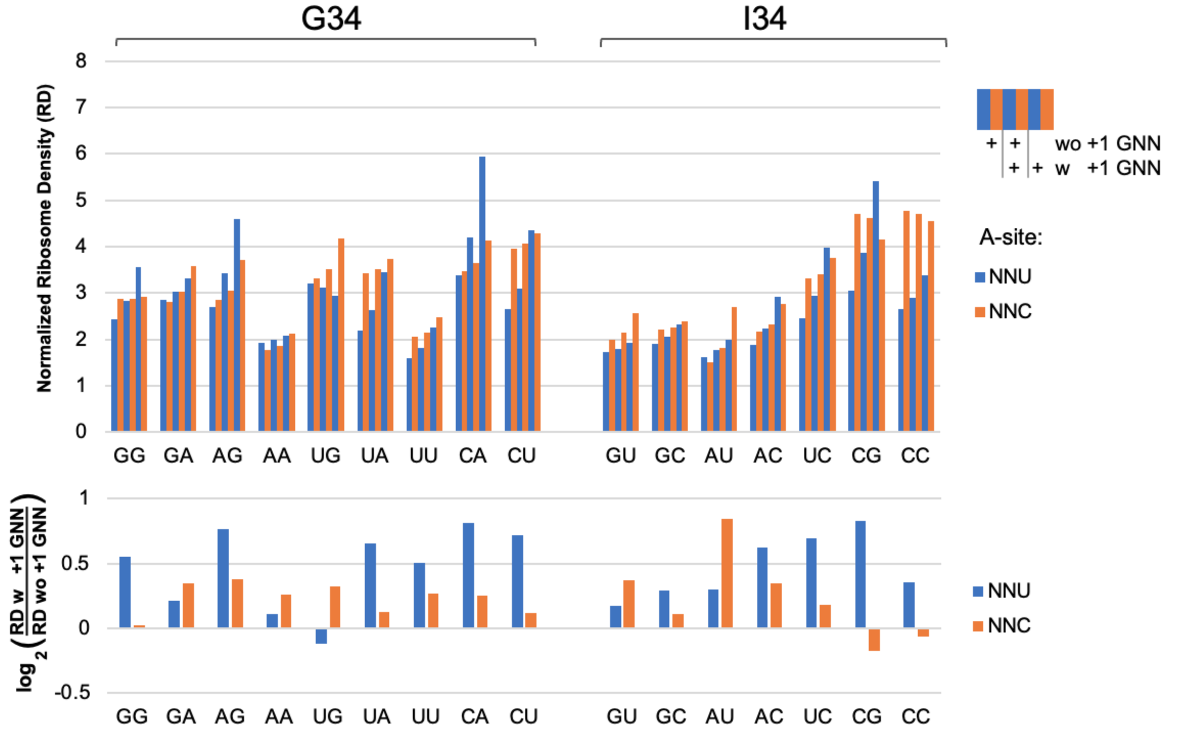


**Fig S6. Pre-accommodation footprints.** Ribosome footprints of length 20-22 nucleotides (21-nt RFPs) are hypothesized to represent pre-accommodation ribosomes that have not accommodated a tRNA at their A-site. The 21-nt RFPs showed similar trends to the larger 28-nt RFPs except that both A-site NNU and NNC codons show higher densities when followed by +1 GNN codon. WT1 and WT2 are independent replicates.

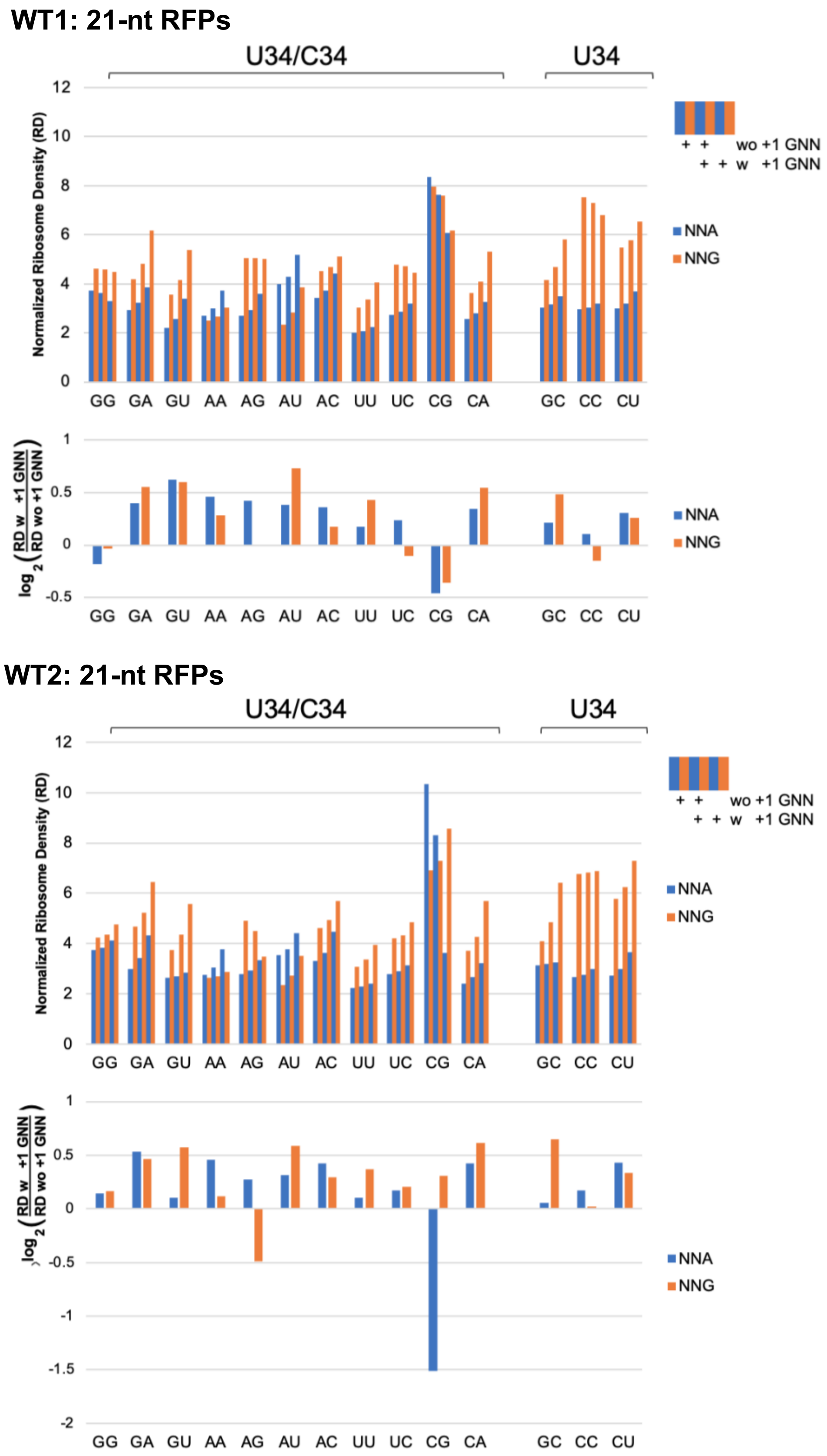


**Fig S7. Pre-accommodation footprints.** Many A-site NNA and NNG codons had elevated ribosome densities when followed by +1 GNN codons (see Table S2). Note that CGA RFPs (replicate WT2) had small numbers of CGA GNN codon pairs with above-threshold ribosome densities (>1 footprint per 10 nt).

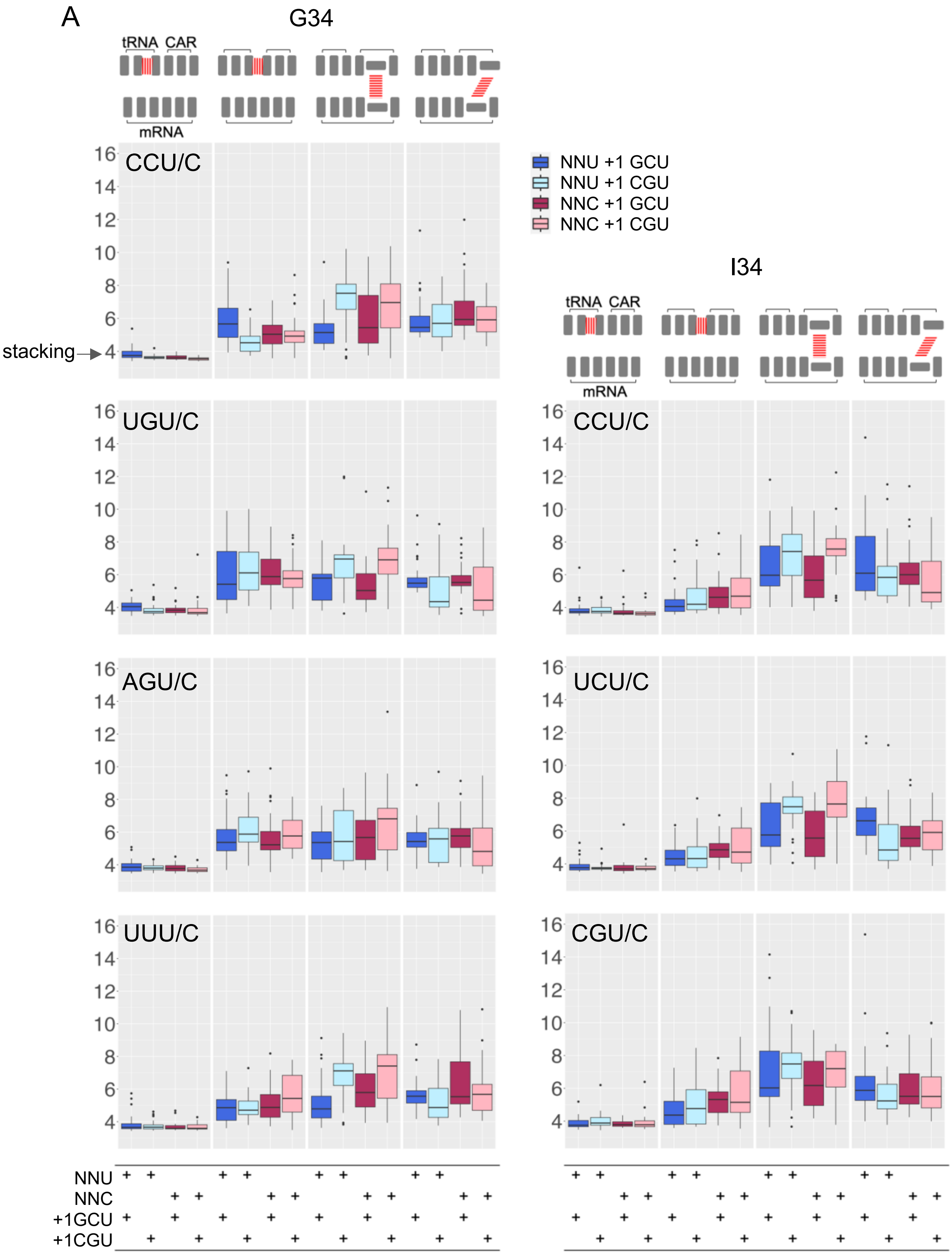
A
G34
tRNA CAR
mRNA
stacking
CCU/C
UGU/C
AGU/C
UUU/C
NNU +1 GCU
NNU +1 CGU
NNC +1 GCU
NNC +1 CGU
I34
tRNA CAR
mRNA
CCU/C
UCU/C
CGU/C
NNU
NNC
+1GCU
+1CGU

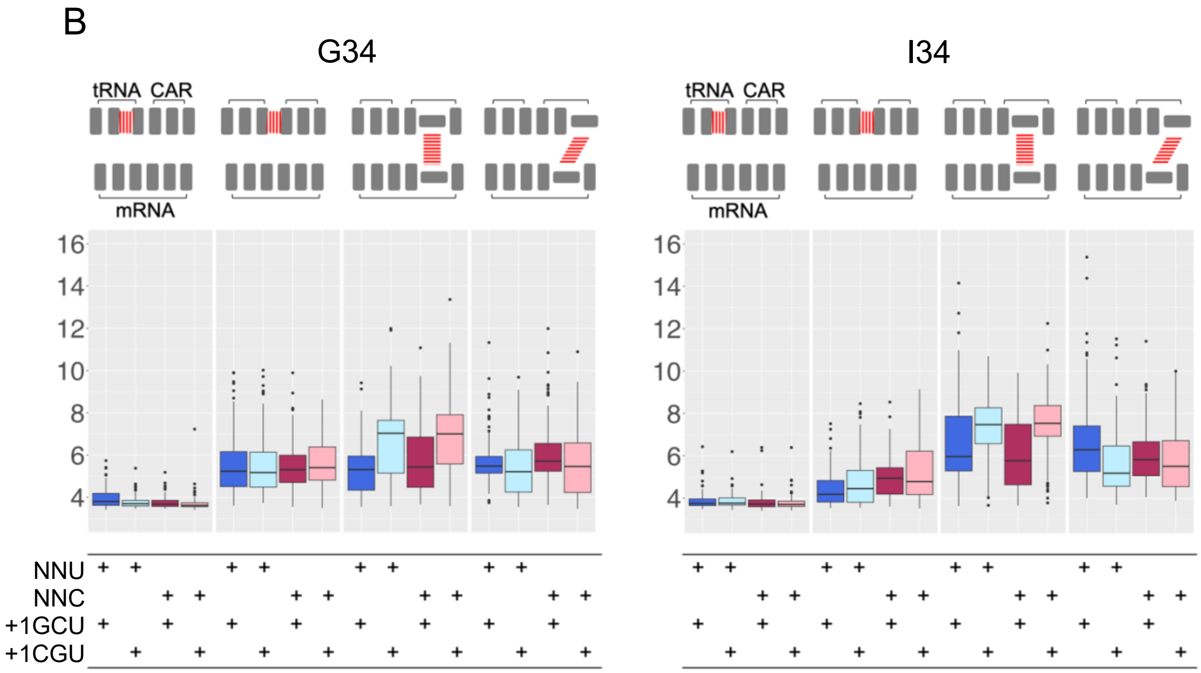


**Fig S8. Stacking behavior of NNU/C codons.** (A) Illustrated left to right (see top diagram) is pi-stacking between bases (tRNA nt35:nt34; nt34:C1054; A1196:+1 nt2) or pi-cation stacking between +1 nt2 and R146 (guanidinium group). Graphs are labeled according to the A-site NNU/C identities, and the NNU or NNC codons are immediately followed by +1 GCU or +1 CGU as summarized in the bottom table. 30 MD replicates were performed for each construct. Stacking was observed when the distances between the centers of geometry of base rings or the guanidinium group of R146 were within 4 Å which due to space constraints resulted in approximately parallel planes of the base rings (or guanidinium group). Nt34:nt35 showed consistent stacking distances of about 4 Å. A-site NNU/C with +1 GCU often showed stacking of A1196 with +1 nt2. A-site NNU/C with +1CGU often showed stacking of R146 with +1 nt2 (see Fig 4). Anchoring of CAR to tRNA nt34 (C1054:nt34) was particularly pronounced for codon pairs (CGU/C, UCU/C, CCU/C) that utilize I34. (B) Collation of data from part A for G34 (120 replicate MD experiments) and I34 (90 replicates) tRNAs.

**Table S1: 28-nt RFPs bootstrap analysis (2-tailed; $p < 0.01$)**

| | +1GNN > wo +1GNN | | +1GNN < wo +1GNN* | |
|---|---|---|---|---|
| | WT1 | WT2 | WT1 | WT2 |
| G34 NNUs | | | | |
| GGU | H | H | - | - |
| GAU | H | H | - | - |
| AGU | H | H | - | - |
| AAU | H | H | - | - |
| UGU | H | H | - | - |
| UAU | H | H | - | - |
| UUU | H | H | - | - |
| CAU | H | H | - | - |
| CUU | H | H | - | - |
| I34 NNUs | | | | |
| GUU | H | H | - | - |
| GCU | H | H | - | - |
| AUU | - | - | L | - |
| ACU | H | H | - | - |
| UCU | H | H | - | - |
| CGU | H | H | - | - |
| CCU | - | H | - | - |
| G34 NNCs | | | | |
| GGC | - | - | - | - |
| GAC | - | - | L | - |
| AGC | - | - | - | - |
| AAC | - | - | L | - |
| UGC | - | - | - | - |
| UAC | - | H | - | - |
| UUC | - | - | - | - |
| CAC | - | - | - | - |
| CUC | - | - | - | - |
| I34 NNCs | | | | |
| GUC | - | - | - | - |
| GCC | - | H | - | - |
| AUC | - | H | - | - |
| ACC | - | - | - | - |
| UCC | - | - | - | - |
| CGC | - | - | - | - |
| CCC | - | - | - | - |

| | +1GNN > wo +1GNN | | +1GNN < wo +1GNN* | |
|---|---|---|---|---|
| | WT1 | WT2 | WT1 | WT2 |
| U34 NNAs | | | | |
| GCA | - | - | L | - |
| CCA | - | - | - | - |
| CUA | - | - | - | - |
| U/C34 NNAs | | | | |
| GGA | - | - | - | - |
| GAA | - | H | - | - |
| GUA | H | H | - | - |
| AAA | - | H | - | - |
| AGA | - | - | - | - |
| AUA | - | - | - | - |
| ACA | - | - | - | - |
| UUA | - | - | - | - |
| UCA | - | - | - | - |
| CGA | - | - | - | - |
| CAA | H | - | - | - |
| U34 NNGs | | | | |
| GCG | H | H | - | - |
| CCG | - | - | - | - |
| CUG | - | H | - | - |
| U/C34 NNGs | | | | |
| GGG | - | - | L | - |
| GAG | H | H | - | - |
| GUG | H | H | - | - |
| AAG | - | - | L | - |
| AGG | - | - | L | L |
| AUG | H | H | - | - |
| ACG | - | H | - | - |
| UUG | H | H | - | - |
| UCG | - | H | - | - |
| CGG | - | H | - | - |
| CAG | - | - | L | L |

* Scored Higher (H) or Lower (L) if A-site codon followed by +1 GNN has significantly higher or lower ribosome densities than when not followed by (wo) +1 GNN (bootstrap 2-tailed analysis, $p < 0.01$). The higher (lower) ribosome density is indicative of slower (faster) translocation rate. Data are for WT1 and WT2 replicate datasets [2].

**Table S2: 21-nt RFPs bootstrap analysis (2-tailed; p < 0.01)**

| | +1GNN > wo +1GNN | | +1GNN < wo +1GNN* | |
|---|---|---|---|---|
| | WT1 | WT2 | WT1 | WT2 |
| G34 NNUs | | | | |
| GGU | H | H | - | - |
| GAU | H | H | - | - |
| AGU | H | H | - | - |
| AAU | - | - | - | - |
| UGU | - | - | - | - |
| UAU | H | H | - | - |
| UUU | H | H | - | - |
| CAU | H | H | - | - |
| CUU | H | H | - | - |
| I34 NNUs | | | | |
| GUU | H | - | - | - |
| GCU | H | H | - | - |
| AUU | H | - | - | - |
| ACU | H | H | - | - |
| UCU | H | H | - | - |
| CGU | H | H | - | - |
| CCU | H | H | - | - |
| G34 NNCs | | | | |
| GGC | - | - | L | - |
| GAC | - | H | - | - |
| AGC | - | - | - | - |
| AAC | H | H | - | - |
| UGC | - | - | - | - |
| UAC | H | - | - | - |
| UUC | - | H | - | - |
| CAC | H | - | - | - |
| CUC | - | - | - | - |
| I34 NNCs | | | | |
| GUC | H | H | - | - |
| GCC | - | - | - | - |
| AUC | H | H | - | - |
| ACC | H | - | - | - |
| UCC | - | - | - | - |
| CGC | - | - | - | - |
| CCC | H | - | - | - |

| | +1GNN > wo +1GNN | | +1GNN < wo +1GNN* | |
|---|---|---|---|---|
| | **WT1** | **WT2** | **WT1** | **WT2** |
| U34 NNAs | | | | |
| GCA | - | - | - | - |
| CCA | - | - | - | - |
| CUA | H | H | - | - |
| U/C34 NNAs | | | | |
| GGA | - | - | - | - |
| GAA | H | H | - | - |
| GUA | H | - | - | - |
| AAA | H | H | - | - |
| AGA | H | H | - | - |
| AUA | H | - | - | - |
| ACA | H | H | - | - |
| UUA | - | - | - | - |
| UCA | - | - | - | - |
| CGA | - | - | - | L |
| CAA | H | H | - | - |
| U34 NNGs | | | | |
| GCG | H | H | - | - |
| CCG | - | - | - | - |
| CUG | H | H | - | - |
| U/C34 NNGs | | | | |
| GGG | - | - | - | - |
| GAG | H | H | - | - |
| GUG | H | H | - | - |
| AAG | H | - | - | - |
| AGG | - | - | - | L |
| AUG | H | H | - | - |
| ACG | - | H | - | - |
| UUG | H | H | - | - |
| UCG | - | - | - | - |
| CGG | - | - | - | - |
| CAG | H | H | - | - |

* Scored Higher (H) or Lower (L) if A-site codon followed by +1 GNN has significantly higher or lower ribosome densities than when not followed by (wo) +1 GNN (bootstrap 2-tailed analysis, $p < 0.01$). The higher (lower) ribosome density is indicative of slower (faster) translocation rate. Data are for WT1 and WT2 replicate datasets [2].

### References for supplemental figures and tables


1. Ghaemmaghami S, Huh WK, Bower K, Howson RW, Belle A, Dephoure N, et al. Global analysis of protein expression in yeast. Nature. 2003;425(6959):737-41. PubMed PMID: 14562106.
2. Wu CC, Zinshteyn B, Wehner KA, Green R. High-Resolution Ribosome Profiling Defines Discrete Ribosome Elongation States and Translational Regulation during Cellular Stress. Mol Cell. 2019;73(5):959-70 e5. Epub 2019/01/29. doi: 10.1016/j.molcel.2018.12.009. PubMed PMID: 30686592; PubMed Central PMCID: PMCPMC6411040.
